# *RIT1*: A Novel Biomarker in Glioblastoma

**DOI:** 10.1101/2020.04.25.061143

**Authors:** Athar Khalil, Georges Nemer

## Abstract

Glioblastoma is the most common type of malignant brain tumors and the most feared cancer among adults. Its high invasiveness and the lack of successful therapies are responsible for the poor prognosis among these patients. A comprehensive understanding for the early molecular mechanisms in glioblastoma would definitely enhance its diagnosis and treatment strategies. Based on in vitro and in vivo models, it was recently postulated that the deregulated expression of key genes that are known to be involved in early neurogenesis could be the instigator of brain tumorigenesis. RIT1 (Ras Like Without CAAX 1) which encodes an unusual “orphan” GTPase protein belongs to this category of critical genes involved in controlling sequential proliferation and differentiation of adult hippocampal neural progenitor cells. As such, we surveyed *RIT1* expression and its potential role in glioblastoma by in-silico means. Our results revealed a significant and progressive upregulation of RIT1 expression in various publicly available datasets. *RIT1* expression ranked among the top upregulated genes in glioblastoma cohorts and correlated with poor overall survival. Genetic and epigenetic analysis of RIT1 didn’t reveal significant aberrations that could underlie its deregulated expression. Moreover, our results pointed on *RIT1* transcriptional activity that could be significantly controlled by STAT3, one of the main players in the onset of glioblastoma. In conclusion, our results are the first proof for the role of RIT1 as an oncogene in glioblastoma paving the way for its potential use as a biomarker and therapeutic target as is the case of the other members in the RAS family.

## Introduction

Brain cancer is a significant source of cancer-related morbidity and mortality worldwide. Although the incidence of neurological cancer is rare accounting for less than 2% of the total number of cancer cases, it is considered among the most feared, aggressive and difficult-to-treat malignancies[1]. Due to their heterogeneity, these tumors are categorized into 29 histologic groups according to the World Health Organization’s (WHO) classification of tumors of the central nervous system (CNS)[2]. Glioblastoma Multiforme (GBM) falls under the category of malignant astrocytoma (grade IV) and is considered the most aggressive and prevalent type of gliomas [3]. GBM is still almost invariably fatal even after aggressive treatment regimens that incorporate neurosurgery, radio- and chemotherapy[4,5]. Several properties make GBM incurable despite the tremendous advancements in basic research and clinical practice. Among these factors are:1- the infiltrative nature of glioma cells making the complete surgical resection impossible even with excessive advancement in neurosurgical techniques, 2-the insufficient delivery of treatments due to pharmacokinetics properties, 3- the blood–brain barrier (BBB), and 4- the developing resistance to conventional chemotherapy and radiation which spares these cells from eradication[2]. Thus, it is imperative to unravel the detailed mechanisms of brain tumorigenesis in order to develop effective and targeted methods for intervention.

RIT1 belongs to the Ras subfamily of GTPases that are known to be implicated in regulating a variety of cellular processes, including cell growth, transformation, differentiation, morphogenesis and apoptosis[6]. Despite harboring a GTPase activity like other proteins from this family, RIT1 lacks the prenylation motif (CAAX, XXCC or CXC) required for its association with the plasma membrane[7]. In humans, its protein is encoded by the *RIT1* gene located on chromosome 1q22 and shares approximately 50% sequence homology with the well-studied *RAS* oncogenes. *RIT1* has a widespread occurrence in the human nervous system and its variants with gain of function mutations are highly associated with Noonan syndrome (NS), an autosomal dominant disorder (OMIM #616564). NS patients develop several partially penetrant developmental anomalies, such as postnatal growth retardation and failure to thrive, congenital heart defects, hypertrophic cardiomyopathy (HCM), hyperkeratosis, and hypotrichosis[8]. Recently, somatic mutations in *RIT1* were shown to be present in several types of cancer including lung adenocarcinoma, hepatoblastoma, urinary tract carcinoma, and adult myeloid malignancies[7]. In mice, *Rit1* is expressed in neurogenic niches of the CNS, the subgranular zone in the dentate gyrus of the hippocampus, and the subventricular zone of the lateral ventricles. Inactivation of *Rit1* doesn’t alter hippocampal development but hippocampal neural cultures derived from *Rit1*^−/−^ mice display increased cell death and blunt MAPK cascade activation in response to oxidative stress[9]. On the contrary, CNS conditional mice model with active RIT1 expression stimulated adult hippocampal neurogenesis and caused an extended neural precursor cells niche[10].

Although the role of *RIT1* in brain development in mammals is well-studied, its expression pattern in human gliomas is still largely unknown. Here, we sought to survey *RIT1* expression in various cohorts and within different grades of human gliomas. We report for the first time a significantly upregulated pattern of this gene in gliomas and suggested a potential role of RIT1 in the progression of the disease. The high expression of *RIT1* was positively associated with decreased survival rate mainly among lower grade glioblastoma patients suggesting its potential prognostic value. Lastly, we demonstrate that *RIT1* expression is positively correlated with that of Signal transducer and activator of transcription 3 (*STAT3)*, the critical mediator of tumorigenesis in GBM, and could be one of its direct targets.

## Material and Methods

### mRNA expression analysis, Kaplan–Meier survival plots and correlation studies for RIT1 in TCGA data

Gene expression profiling interactive tool (GEPIA; http://gepia.cancer-pku.cn/index.html) is an interactive web server that uses a standard processing pipeline for analyzing the RNA sequencing expression data of 9,736 tumors and 8,587 normal samples from the TCGA and the GTEx projects[11]. Boxplots were performed to visualize the expression of RIT1 in various tumors versus their matched normal tissues. For analysis we used ANOVA statistical method for differential gene expression analysis, log2(TPM + 1) transformed expression data for plotting, TCGA tumors compared to TCGA normal and GTEx normal for matched normal data, |log2FC| cutoff of 1, and a q-value cutoff of 0.01. Differential gene studies and survival analysis based on RIT1 or STAT3 mRNA level in lower grade glioma(LGG; Tumor: n=518; Normal: n=207) and glioblastoma multiforme (GBM Tumor: n=163; Normal: n=207) data in TCGA was carried on. The overall survival with a median group cutoff was displayed on the plot along with the hazard ratio (HR) and log rank P value. The correlation between the expression of *RIT1* and *STAT3* in LGG and GBM was determined using Spearman’s correlation analysis.

### *RIT1* expression analysis in various glioma datasets

We used the Oncomine webportal to analyze *RIT1* mRNA expression in various human glioma cohorts. Oncomine (https://www.oncomine.org/resource/login.html) is an online database consisting of previously published and open-access microarray data englobing 715 datasets and 86,733 samples[12]. The mRNA expression of *RIT1* in clinical brain cancer tissue was compared with that in normal control, using a Students’ t-test. The search filters were set up as RIT1 for gene name, cancer vs normal analysis, and Brain/CNS cancer for cancer type. Thresholds were set as follows: gene rank, 10%; fold change, 2; and P-value, 0.01. Analysis were done on TCGA, GSE2223, GSE4290, GSE7696 and unavailable GEOdataset by Shai et.al [13]. Using Genevestigator v3 suite we evaluated *RIT1* expression among 632 cancer categories from data selection. Level of expression of *RIT1* based on the signal intensity on Affymetrix Human Genome U133 Plus 2.0 Array was displayed. Then we extracted samples with the highest 5% expression of *RIT1* amongst all categories.

### Genetic and epigenetic alterations of RIT1 in glioblastoma

The cBioPortal for Cancer genomics is an open-access resource (http://www.cbioportal.org/), providing visualization and analyzing tool for more than 5,000 tumor samples from 105 cancer studies in TCGA pipeline. The term “RIT1” was searched within the cBioPortal database and a cross-cancer summary was obtained for it. The search parameters included alterations (amplification, deep deletion, missense mutations), copy-number variance (CNV) from GISTIC and RNA seq data with the default setting. In order to see the effect of epigenetic modification i.e. DNA methylation, we analyzed the methylation pattern of *RIT1* in promoter region (mainly CpG islands) of normal vs glioblastoma specimens using the Wanderer web tool (http://maplab.imppc.org/wanderer/doc.html) [14].

### Analysis of the RIT1 gene for promoter regulation by STAT3

We used the **INSECT**2.0 (**IN**-silico **SE**arch for **C**o-occurring **T**ranscription factors, version 2.0) web server for prediction and analysis of the human *RIT1* promoter for potential binding sites for the transcription factor STAT3 (https://bioinformatics.ibioba-mpsp-conicet.gov.ar/INSECT2/index.php). The chosen binding sites for STAT3 were based on the Jaspar (Murine) and Transfac (Human) databases, and the analysis include a phylogenetic assessment of the corresponding murine *RIT1* promoter.

## Results

### Differential expression studies of RIT1 transcript in pan-cancer

We surveyed the expression pattern of *RIT1* amongst different cancer types by *in silico* analysis of publicly available expression datasets within the gene expression profiling interactive tool (GEPIA) project constituting RNA sequencing data from 9,736 tumors and 8,587 normal samples. Our analysis demonstrated that *RIT1* expression was significantly upregulated only in four cancer subtypes amongst which are Glioblastoma Multiforme (GBM) and Lower Grade Glioma (LGG) (Figure-1A). Studying *RIT1* expression in 632 different categories of caner that are presented in Genevestigator database revealed that those with the highest expression correspond mainly to brain glioma samples (Supplementary Figure-1). Among the 5% categories that scored the highest RIT1 expression; 25% showed to be corresponding to gliomas. Furthermore, mRNA expression from GEPIA online database showed that *RIT1* is upregulated by 2 and 2.5-log2 fold change in LGG and GBM, respectively, as compared to matched normal tissues retrieved from TCGA and Gtex (Figure-1B).

**Figure 1:**
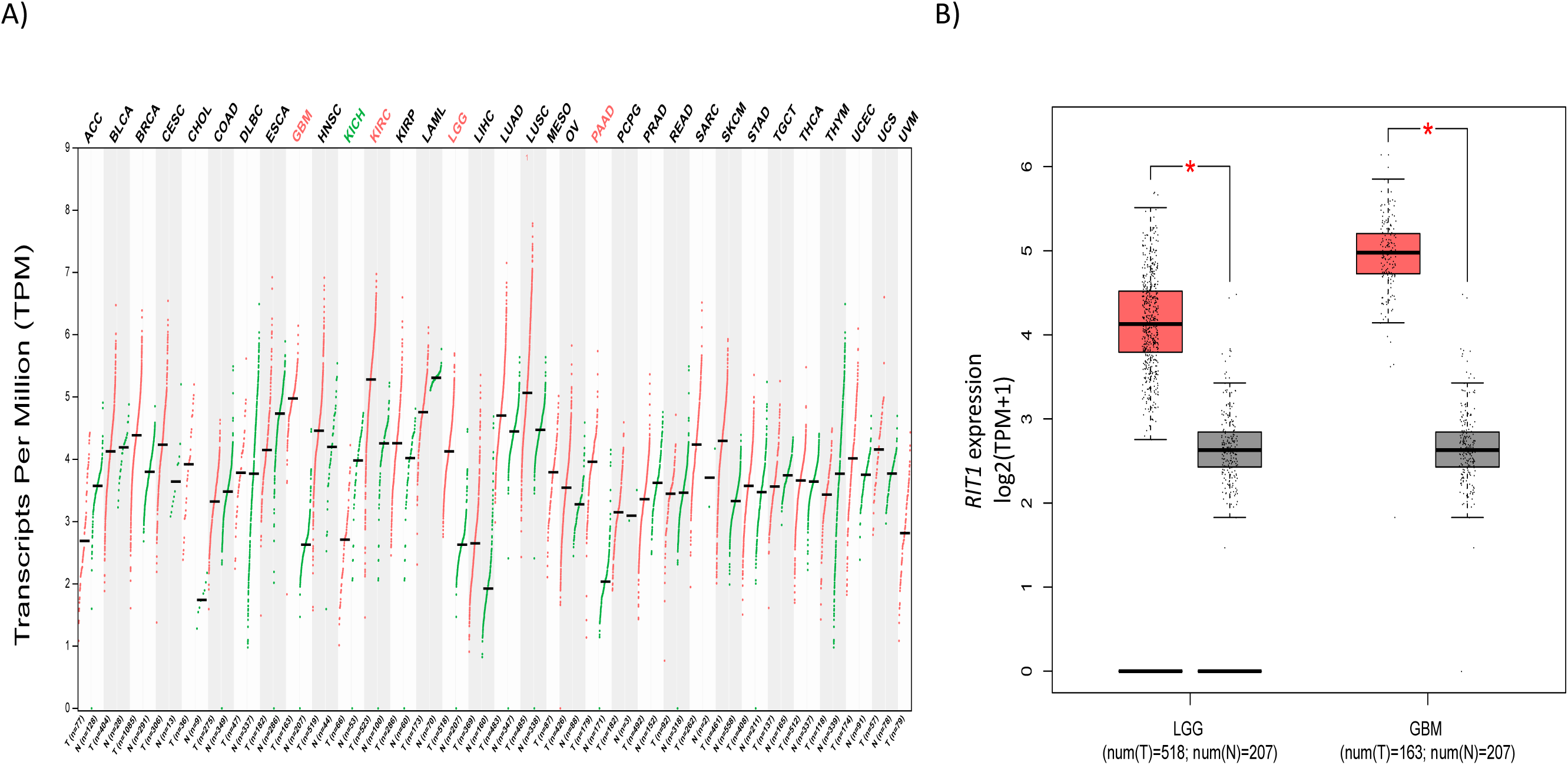
Expression profile for *RIT1* across cancer types. A) Expression levels for *RIT1* across 33 TCGA tumors compared to TCGA normal and Genotype-Tissue Expression (GTEx) data using the GEPIA webserver. For each TCGA tumor type (red), its matched normal and GTEx data (green) are given; T: tumor; N: normal; n: number. Y axis: transcript per million (log2(TPM + 1)). X axis: number of tumor and normal samples. ACC: adrenocortical carcinoma; BLCA: bladder urothelial carcinoma; BRCA: breast invasive carcinoma; COAD: colon adenocarcinoma; CHOL: cholangio carcinoma; CESC: cervical squamous cell carcinoma and endocervical adenocarcinoma; DLBC: lymphoid neoplasm diffuse large B-cell lymphoma; ESCA: esophageal carcinoma; GBM: glioblastoma multiforme; HNSC: head and neck squamous cell carcinoma; KICH: kidney chromophobe; KIRC: kidney renal clear cell carcinoma; KIRP: kidney renal papillary cell carcinoma; LAML: acute myeloid leukemia; LGG: brain lower grade glioma; LIHC: liver hepatocellular carcinoma; LUAD: lung adenocarcinoma; LUSC: lung squamous cell carcinoma; MESO: mesothelioma; OV: ovarian serous cystadenocarcinoma; PCPG: pheochromocytoma and paraganglioma; PAAD: pancreatic adenocarcinoma; PRAD: prostate adenocarcinoma; READ: rectum adenocarcinoma;SARS: sarcoma; SKCM: skin cutaneous melanoma; STAD: stomach adenocarcinoma; TGCT: testicular germ cell tumors; THCA: thyroid carcinoma; THYM: thymoma; UCEC: uterine corpus endometrial carcinoma; UCS: uterine carcinosarcoma; UVM: uveal melanoma. B) Expression level for *RIT1* in LGG and GBM in TCGA versus matching normal TCGA tissues and Gtex data presented as log2(TPM+1) scale (*p < 0.01). Tumor group: red column; non-tumor group: gray column.

### Upregulated expression of RIT1 in human gliomas relative to normal brain tissues

To further explore the differential expression of *RIT1* between glioblastoma and non-cancerous brain tissues, the TCGA data as well as three other glioblastoma cohorts were analyzed using the Oncomine online database web portal (Figure-2A). All four datasets showed a significant increase in the expression of *RIT1* in glioblastoma tissues compared to that of normal brain (p<0.01; Figure-2B). *RIT1* scored among the top 4% differentially upregulated genes in the TCGA-GBM dataset. This high score was also identified in additional datasets where *RIT1* scored among the highest differentially upregulated genes in GBM tissues (7% in GSE2223 and GSE4290, 2% in GSE7696)[15-17]. These results pointed to a commonly and significant consistent overexpression of the *RIT1* in human glioblastoma.

**Figure-2:**
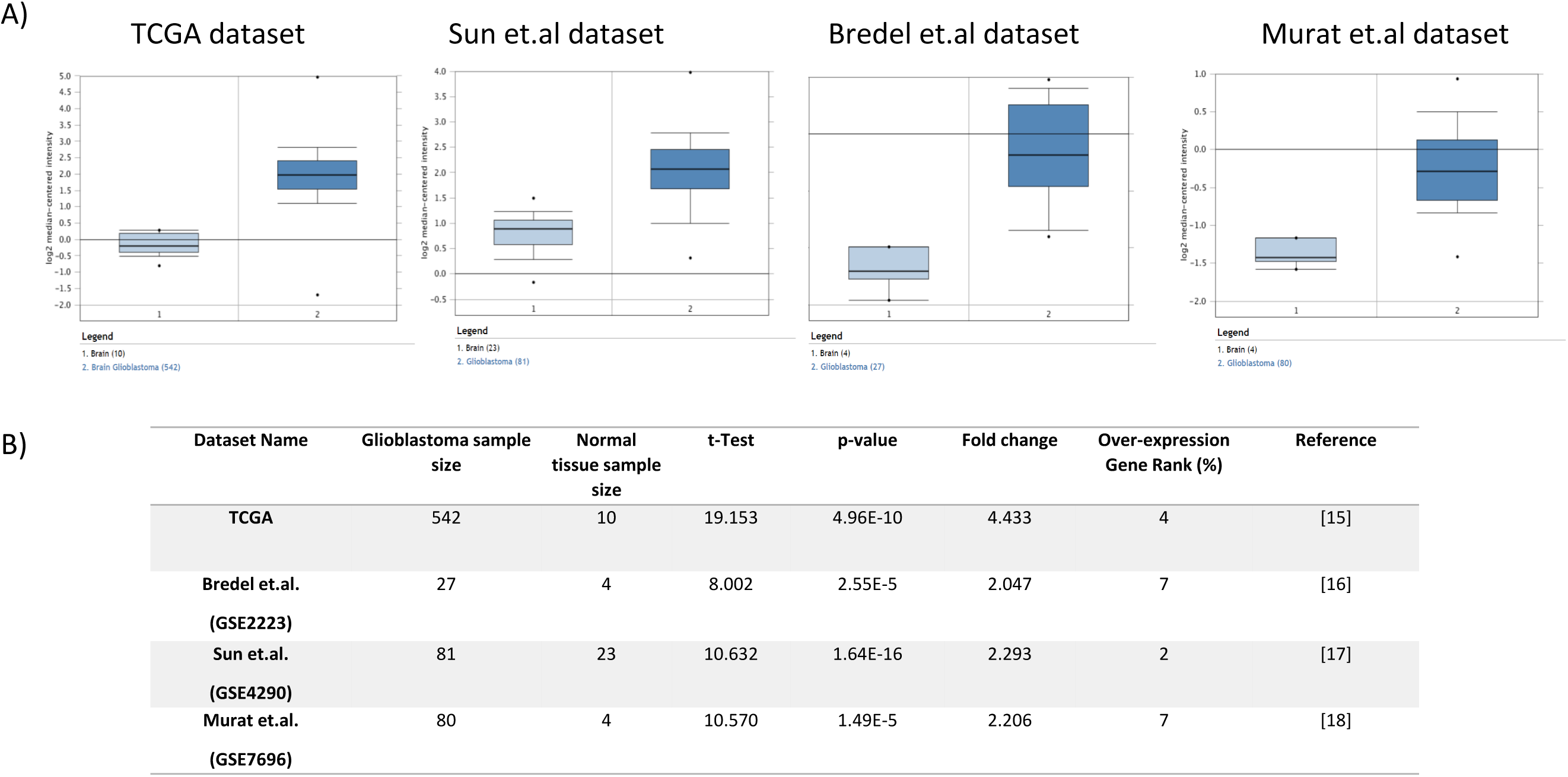
*RIT1* analysis in different glioblastoma cohorts. Boxplots comparing RIT1 expression in normal (left plot) and cancer tissue (right plot) that were derived from Oncomine database (A). The fold change of RIT1 in various glioblastoma datasets presented (B) expressed as boxplots for the TCGA, Bredel et.al, Sun et.al, and Murat et.al datasets. X-axis of the plot represents normal vs cancer group; Y-axis represents mRNA expression in log2 median/mean centered intensity. The line in the middle represents the median value. Differences were examined statistically by two-tailed student’s t-test.

### A progressive upregulation of RIT1 and its potential prognostic value

Global transcriptional profiles of gliomas of different types and grades are distinct from each other and from that of normal brain. Thus, we studied the expression of *RIT1* from Shai.et.al dataset in varying pathologic types and grades of gliomas (low-grade astrocytomas (grade II), the anaplastic astrocytomas and oligodendrogliomas (grade III) and the malignant glioblastoma (grade IV)) as well as seven normal brain samples that were taken from the subcortical white matter[13]. The observed up-regulated pattern of *RIT1* was progressive as we move from lowest grade to more malignant form of glioblastoma (Figure-3A). These findings suggest that the up-regulation of the *RIT1* is possibly implicated in the development of premalignant precursors and in the progression of these lesions to malignant glioblastoma. Additionally, to evaluate the possible prognostic value for *RIT1* in different grades of glioblastoma we performed Kaplan-Meier survival analysis according to the median mRNA expression of *RIT1* in both GBM and LGG using Gepia analysis tool. The results showed that high *RIT1* mRNA expression (HR =5.7, 95% CI) was associated with worse overall survival for glioblastoma patients but the results didn’t reach statistical significance (log-rank P=0) (Figure-3B). While when *RIT1* evaluation was separately done on GBM and LGG patients, only the latter showed a statistically significant association between high *RIT1* expression and poor overall survival (Figure-4A,B). Therefore, a high *RIT1* expression score might be associated with decreased survival time mainly among lower grade glioma patients.

**Figure-3:**
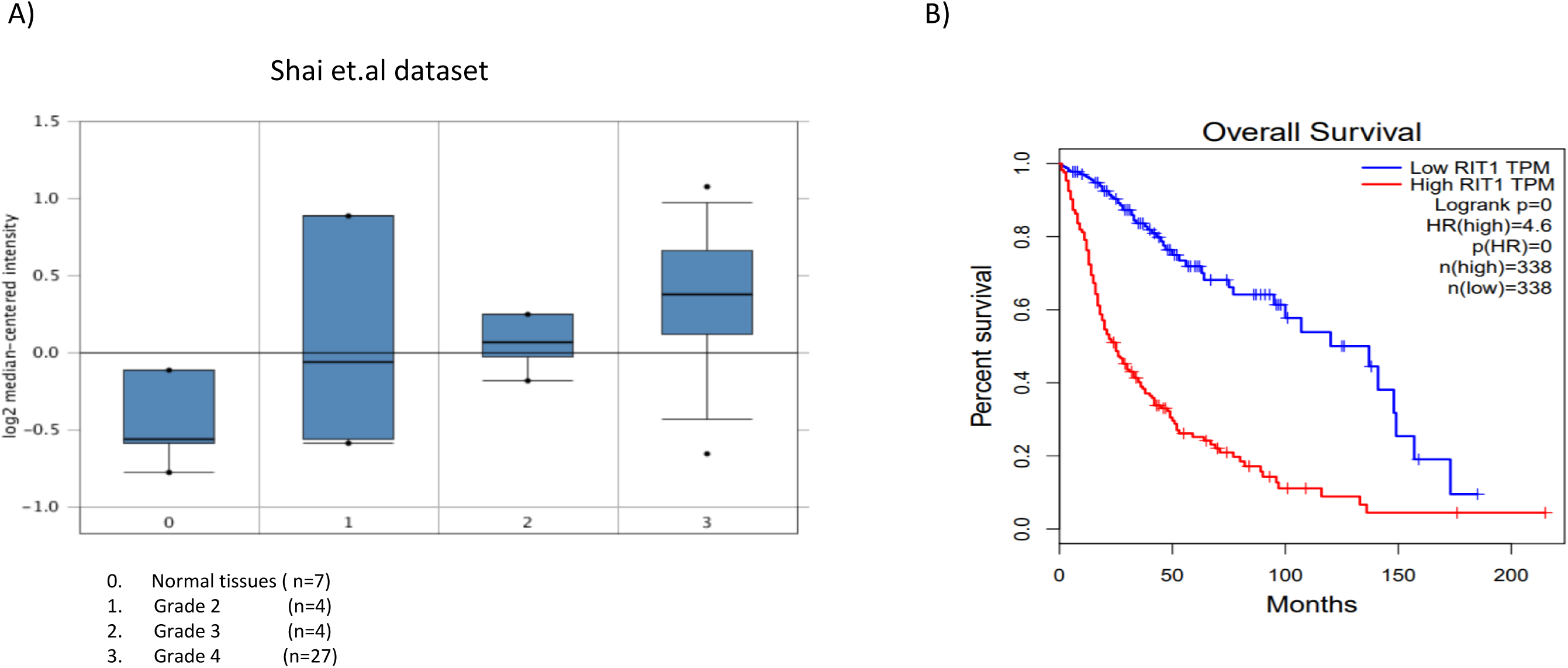
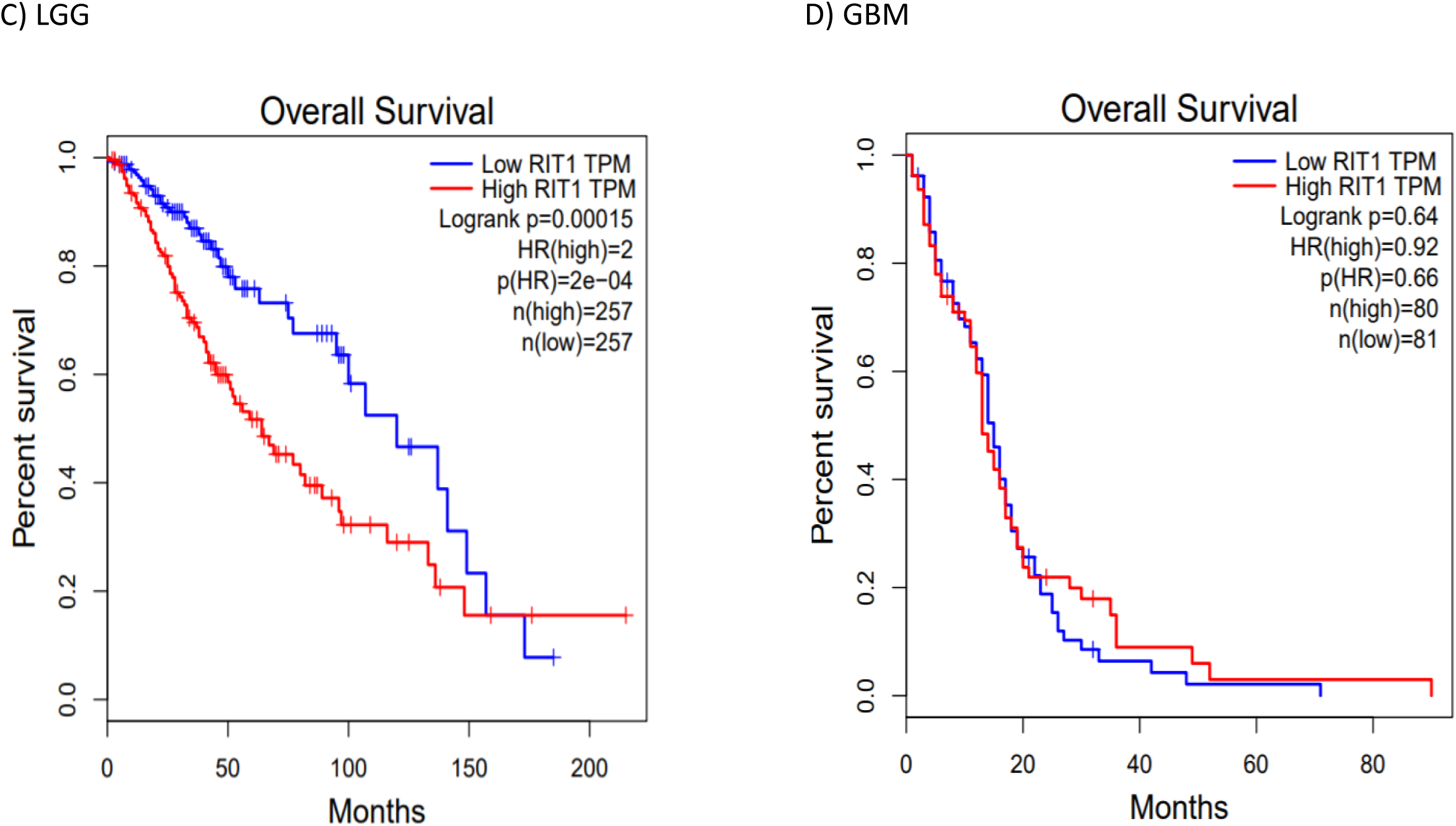
Association of *RIT1* expression levels with three glioma grades and overall survival time. A) RIT1 expression stratified by glioblastoma grade versus normal tissues in the Shai glioblastoma dataset derived from Oncomine software. B) Kaplan–Meier plots of survival were generated by Gepia (http://gepia.cancer-pku.cn) using the data from TCGA comparing overall survival of patients according to *RIT1* expression in LGG and GBM together. Red line indicates the samples with gene highly expressed, and blue line shows the samples with gene lowly expressed. HR, hazard ratio.

**Figure-4:**
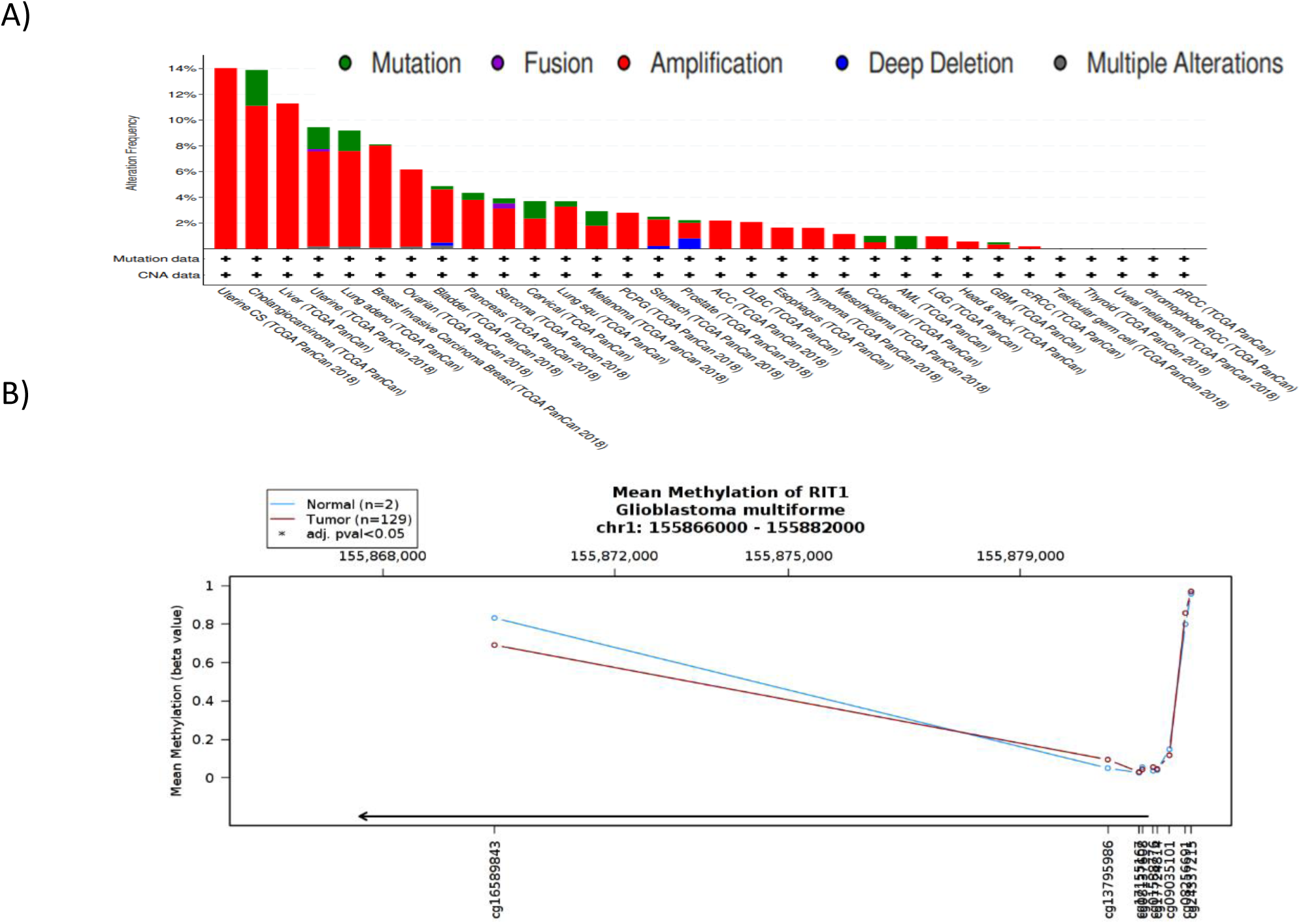
Low *RIT1* expression is better correlated with survival in LGG. Kaplan–Meier plots of survival were generated by Gepia (http://gepia.cancer-pku.cn) using the data from TCGA comparing overall survival of patients according to *RIT1* expression in C) LGG or D) GBM versus matched normal tissues from Gtex. Red line indicates the samples with gene highly expressed, ad blue line shows the samples with gene lowly expressed. HR, hazard ratio.

### Unrevealed mechanism of action that might underlie the upregulation of RIT1 in GBM

Then we sought to understand the molecular mechanism responsible for upregulating *RIT1* expression in various glioblastoma stages. For this reason, we surveyed *RIT1* alteration frequency of mutations and copy number alterations (CNAs) in all types of cancer using cBioportal database. Our analysis revealed only a minor contribution (0.6% of the cases) of such alterations in glioblastoma (Figure-5A). We then opted to investigate the methylation status of the promoters of *RIT1* gene using TCGA Wanderer, an interactive web viewer for the visualization of DNA methylation based on TCGA data. The results showed no significant change in the methylation pattern of *RIT1* promoter between glioblastoma and normal brain tissues which highlight the need for understanding the mechanism(s) that underlie the observed upregulated pattern of *RIT1* in glioblastoma (Figure-5B).

**Figure-5:**
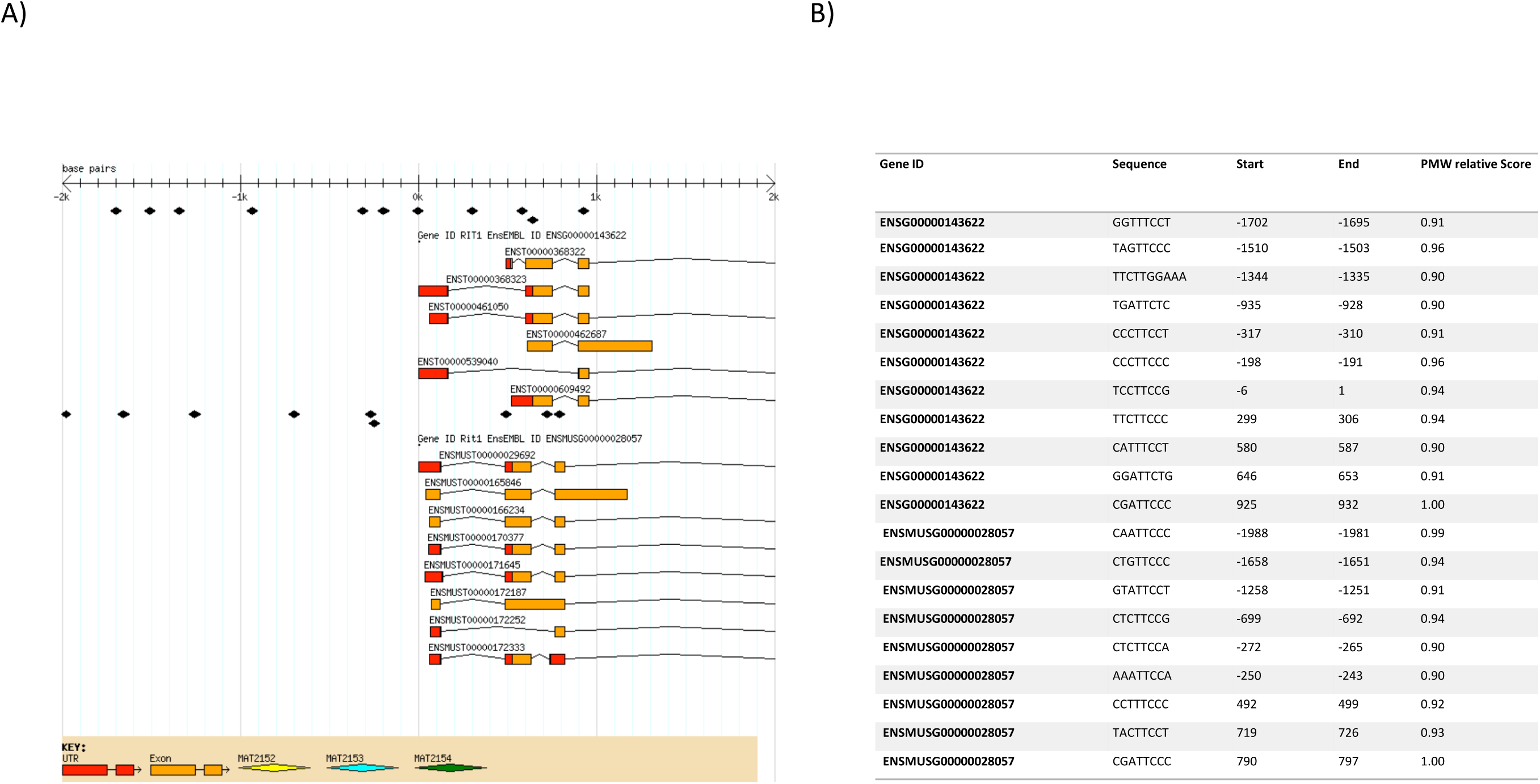
Genetic aberrations of *RIT1* in cancer subtypes and its DNA methylation status in GBM. A) The alteration frequency of a *RIT*1 gene signature was determined using the cBioPortal (http://www.cbioportal.org). The alteration frequency included deep deletions (blue), amplification (red), multiple alterations (grey) or mutation (green). B) Graphics obtained from Wanderer data base (http://maplab.imppc.org/wanderer/) of all CpGs island of *RIT1* and its percentage of DNA methylation in GBM (medium of 129 samples, Red) and normal brain tissues (medium of 2 samples, blue). In x-axis the cg# indicates the position of the CpG island. *: indicates differences statistically significant between normal and tumor samples

### STAT3 a potential transcriptional regulator for RIT1 in glioblastoma

We used the INSECT2.0 software to assess the potential transcriptional regulation of RIT1 by STAT3 a well-established oncogenic driver of GBM. Based on TRANSFAC and JASPAR databases, we searched for potential binding sites over the upstream 2000bp promoter and 1000bp downstream regions (relative to the transcription start site) on the *RIT1* human gene and its ortholog in mice. We set the position weight matrix (PWM) at >90% which revealed a total of 11 potential binding sites, two of them with an overlapping pattern in mice (Figure-6A,B). To support our finding, we showed a significant positive correlation between RIT1 and STAT3 mRNAs, both in glioblastoma(R=0.7, p=1.9e-100) and normal (R=0.81, p=5.6e-72) tissues (Cerebellum, Cortex and Hippocampus) where RIT1 is known to be highly expressed (Figure-7A,B)[18]. *STAT3* expression level was further analyzed in LGG and GBM versus normal matched TCGA and GTEx data. Consistent with previous studies, we showed a significant upregulation of *STAT3* in both grades of gliomas with a higher expression in GBM (Supplementary Figure-2). Further investigations and in-vitro studies are warranted to confirm a direct transcriptional regulation of *RIT1* by *STAT3*.

**Figure-6:**
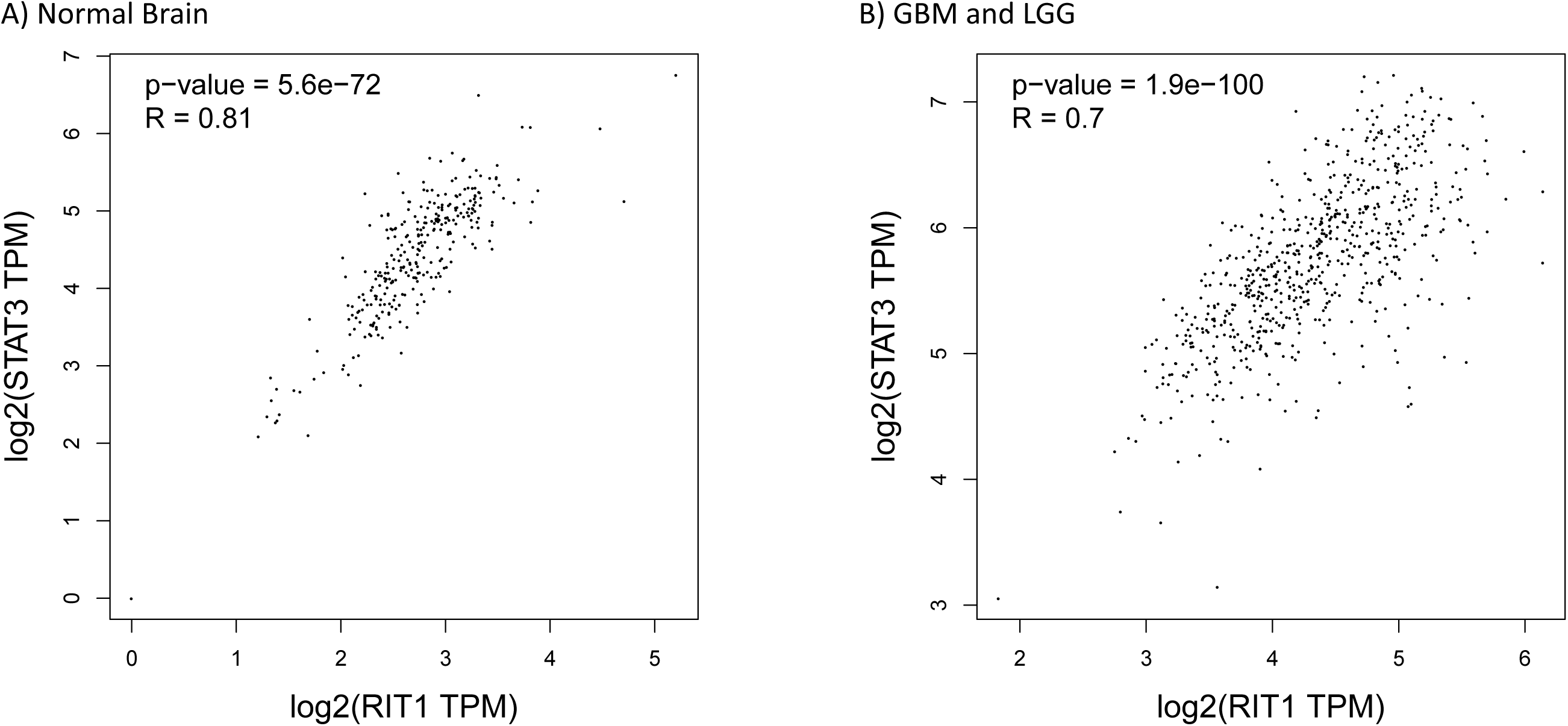
In silico analysis of the *RIT1* promoter for potential *STAT3* binding sites. The STAT3 potential binding sites with their corresponding position (Start, End) on the human (ENSG00000143622) and murine (ENSMUS00000028057) Rit1 gene (A), are visualized over the intronic/exonic junctions (B). Negative numbers indicate positions in the promoter region whereas positive numbers indicate regions downstream the transcriptional starting site. PWM: position weight matrix.

**Figure-7:**
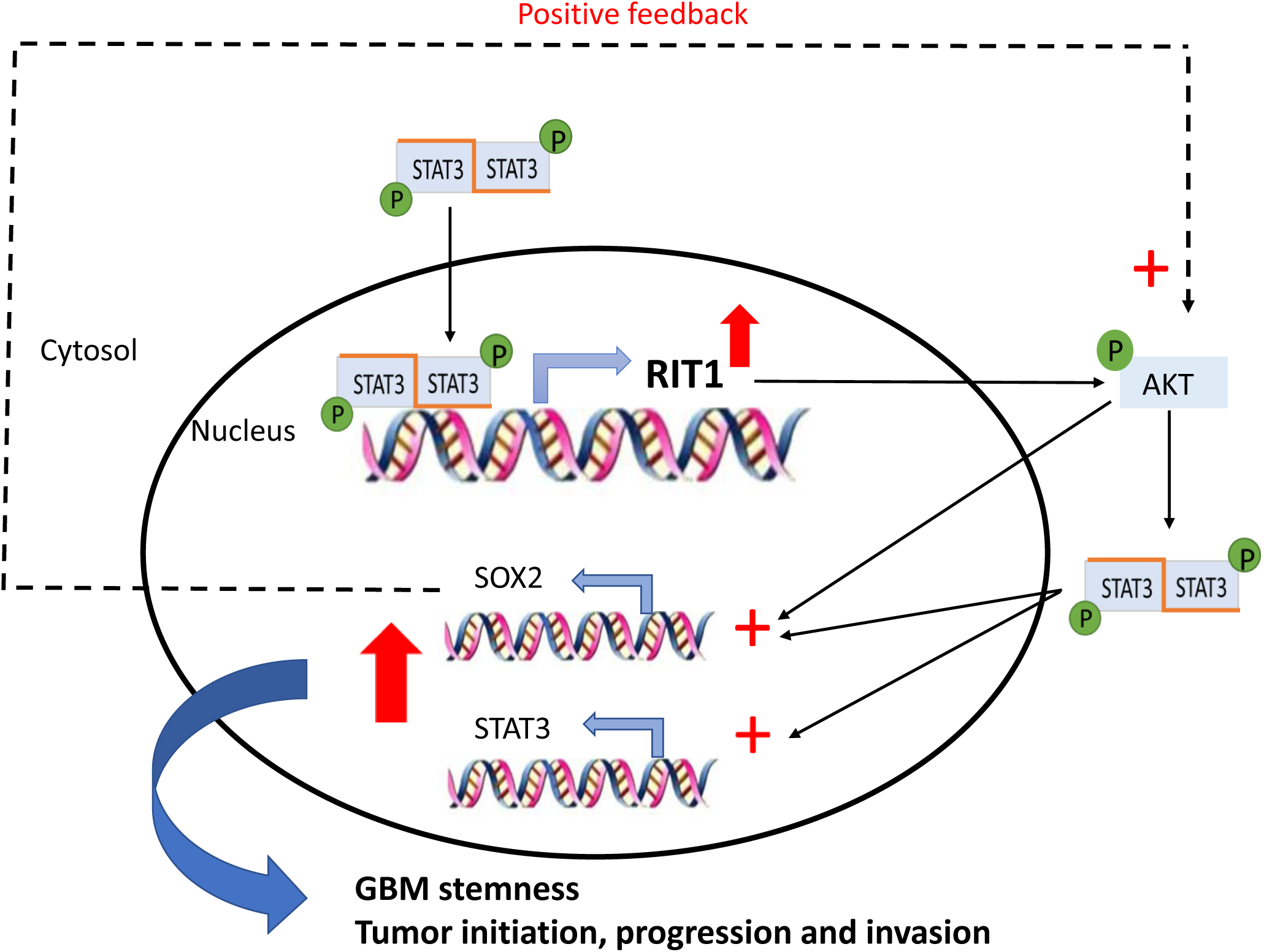
Spearman correlation analysis of *RIT1* and *STAT3* gene and mRNA expression levels. The expression of RIT1 was positively correlated with STAT3 mRNA expression in A) normal brain tissues (r=0.81, P<0.001) and B) glioblastoma tissues (r=0.7, P<0.001).

## Discussion

The present study aimed at investigating the expression and the prognostic value of *RIT1* in glioblastoma using integrated large databases. Previous *in-silico* and experimental studies on *RIT1* have documented its role in numerous cancer types and in the neurogenesis process, yet its expression and prognostic value in GBM was not assessed. We found that *RIT1* mRNA expression level in GBM tissues was significantly higher than that in non-tumor tissues (p< 0.001). Moreover, the upregulation of *RIT1* expression followed a progressive pattern as we move from lower grade glioma to the malignant multiforme. In contrast, *RIT1* did not tend to be altered in the GBM genomic signature, and its methylation pattern was not significantly disrupted. The higher expression of *RIT1* was significantly correlated with poor prognostic survival outcome, mainly in LGG. Finally, we proposed a novel molecular insight in the genetic network controlling glioblastoma by linking *RIT1* overexpression to the well-documented oncogenic player, *STAT3*.

Using transcriptome sequencing, we demonstrated for the first time a significant upregulated expression of *RIT1* in glioma that scored among the top upregulated genes in various GBM cohorts. The mechanism responsible for deregulating *RIT1* expression in GBM is still poorly understood. Previous studies have shown that gain of chromosome 1q, particularly in the region of 1q21-25, is the most frequent finding (29%) among childhood high grade gliomas. This is even accentuated with the finding of a significant association between 1q gain and decreased survival among these patients. Thus the authors proposed the need for the identification of one or more candidate genes specifically on that region to be the key for identifying pathways involved in the biology of these tumors and as reliable prognostic biomarker(s)[19]. It is noteworthy to mention that *RIT1* maps to 1q22 suggesting that the upregulated expression of this gene in both LGG and GBM might simply be a surrogate of the copy number gain of chromosome 1q. Forthcoming work is warranted to validate the expression of the *RIT1* subfamily in LLG and GBM using orthogonal measures (e.g., immunohistochemistry). Other potential mechanisms responsible for *RIT1* overexpression could be hypomethylation, promoter/enhancer activation by highly expressed TFs, or downregulation of miRNAs that would normally inhibit *RIT1* expression.

The frequency of genetic alterations (mutations, CNVs, deletions…etc.) in *RIT1* within GBM was not high enough (≃1%) to explain its consistent upregulation among all studied cohorts. Methylation was ruled out, but further studies should be carried-on due to the lack of representable number of normal brain tissues in the methylation experiments. miRNAs expression profile would only be beneficial if there are experimental proofs on the direct regulation of *RIT1*. Based on the miRTarBase which provides information about experimentally validated miRNA-target interactions (MTIs), three human miRNAs were predicted to directly target *RIT1* (miR-125a-5p, miR-218-5p and miR-340-5p[20]. The three were shown to be significantly downregulated in GBM, and their role in tumor initiation, progression and invasion was well-defined[21-23]. Yet, these studies pointed out to distinct targeted genes regulated by each miRNA while neglecting their common target; *RIT1*. Further studies will be needed to evaluate the impact of the downregulation of those miRNAs specifically on *RIT1* expression and consolidate the hypothesis about their common target in GBM tumorigenesis.

Lastly, our promoter analysis revealed a couple of binding sites that could be implicated in regulating transcription of *RIT1*. Amongst these transcriptional factors is *STAT3*, a member of the STAT family of transcription factors, known to drive several pro-oncogenic mechanisms that promote cell proliferation, survival, immune suppression, invasion, and angiogenesis in GBM [24]. The consecutive activation of *STAT3* resulting from either gain-of-function mutations or overexpression of upstream growth factor receptors is detected in 90% of human GBM tumors and GBM cell lines[25]. Moreover, the hypomethylated pattern of *STAT3* in the acquired recurrent glioma-CpG island methylator phenotype (G-CIMP) is characterized with unfavorable overall survival [26]. Thus, by combining the detected activation of STAT3 with it potential transcriptional binding activity to *RIT1* promoter, one could explain partially the upregulated pattern of *RIT1* in glioblastoma. Ultimately, in-vitro studies are warranted to confirm *RIT1* as a novel direct downstream target for STAT3.

In parallel, we could also hypothesize that RIT1 can modulate directly or indirectly *STAT3* expression and/or transcriptional activity. This hypothesis is based on the fact that the primary target for *RIT1*-kinase function is the serine/threonine kinase (AKT) which is in turn a key player in stimulating the transcriptional activity of both STAT3 and SOX2 [10]. SOX2, a member of the Sex-determining region Y (SRY)-box (SOX) factors, is considered as the most enriched gene among the stemness signature in GBM stem cells (GSCs)[30]. This transcriptional factor is overexpressed in malignant glioma from several different and independent cohorts while displaying minimal expression in normal tissues[29]. Additionally, in-vitro and in-vivo studies confirmed that RIT1 activation can enhance hippocampal neural progenitor cells proliferation through activating SOX2 transcriptional activity in an Akt-dependent manner[10]. Thus, *RIT1* aberrant expression and its positive correlation with *SOX2* pinpoint to the critical role played by *RIT1* in maintaining the self-renewing properties of GBM stem cells. Taken into consideration that SOX2 was recently acknowledged to regulate p-AKT and p-STAT3 in GBM, we could come up with a scenario of three main players RIT1, STAT3, and SOX2 that can commit a positive feedback loop in favor for GBM tumorigenesis (Figure-8).

**Figure-8:** Schematic model of the *STAT3*-*RIT1*-*SOX2* pathway in glioblastoma. p-STAT3 is a well-documented upregulated active protein in GBM that could enhance *RIT1* transcription which in turn upregulates the expression of *STAT3* and *SOX2* in an AKT dependent manner. SOX2 could exert a positive feedback promoting phosphorylation of AKT and STAT3. The activation of this loop and the upregulation of *STAT3* and *SOX2* will enhance GBM stemness and the tumorigenesis process.

## Conclusion

This study presented for the first time *RIT1* gene encoding a RAS protein, as an oncogene in glioblastoma and opened the door for further investigation on its potential use as therapeutic target and molecular biomarker. The RAS subfamily of GTPases created an evolution in the field of cancer. Members of this family are among the most studied oncogenes and are used in the molecular classification of various tumors. Adding *RIT1* to this list, doesn’t only expand our knowledge in the field, but would have a major impact on the health system if targeted by specific inhibitors to successfully cure glioblastoma and/or other genetic diseases where it’s overexpressed like in Noonan syndrome.

## Acknowledgements

Not applicable.

## Figures Legends

**Supplementary Figure-1:** *RIT1* expression among 632 cancer categories analyzed using GENEVESTIGATOR.

**Supplementary Figure-2: Expression** level of *STAT3* in LGG and GBM versus matching normal TCGA tissues and Gtex data presented as log2(TPM+1) scale (*p < 0.01). Tumor group: red column; non-tumor group: gray column. TPM, transcripts per kilobase million.

## References

1. de Robles P, Fiest KM, Frolkis AD, Pringsheim T, Atta C, St Germaine-Smith C, Day L, Lam D, Jette N (2015) The worldwide incidence and prevalence of primary brain tumors: a systematic review and meta-analysis. Neuro Oncol 17 (6):776–783. doi: 10.1093/neuonc/nou283

2. Ostrom QT, Gittleman H, Fulop J, Liu M, Blanda R, Kromer C, Wolinsky Y, Kruchko C, Barnholtz-Sloan JS (2015) CBTRUS Statistical Report: Primary Brain and Central Nervous System Tumors Diagnosed in the United States in 2008-2012. Neuro Oncol 17 Suppl 4:iv1–iv62. doi: 10.1093/neuonc/nov189

3. Tamimi AF, Juweid M (2017) Epidemiology and Outcome of Glioblastoma. In: De Vleeschouwer S (ed) Glioblastoma. Brisbane (AU). doi: 10.15586/codon.glioblastoma.2017.ch8

4. Azzarelli R, Simons BD, Philpott A (2018) The developmental origin of brain tumours: a cellular and molecular framework. Development 145 (10). doi: 10.1242/dev.162693

5. Gomes AL, Reis-Filho JS, Lopes JM, Martinho O, Lambros MB, Martins A, Schmitt F, Pardal F, Reis RM (2007) Molecular alterations of KIT oncogene in gliomas. Cell Oncol 29 (5):399–408

6. Lee CH, Della NG, Chew CE, Zack DJ (1996) Rin, a neuron-specific and calmodulin-binding small G-protein, and Rit define a novel subfamily of ras proteins. J Neurosci 16 (21):6784–6794

7. Cave H, Caye A, Ghedira N, Capri Y, Pouvreau N, Fillot N, Trimouille A, Vignal C, Fenneteau O, Alembik Y, Alessandri JL, Blanchet P, Boute O, Bouvagnet P, David A, Dieux Coeslier A, Doray B, Dulac O, Drouin-Garraud V, Gerard M, Heron D, Isidor B, Lacombe D, Lyonnet S, Perrin L, Rio M, Roume J, Sauvion S, Toutain A, Vincent-Delorme C, Willems M, Baumann C, Verloes A (2016) Mutations in RIT1 cause Noonan syndrome with possible juvenile myelomonocytic leukemia but are not involved in acute lymphoblastic leukemia. Eur J Hum Genet 24 (8):1124–1131. doi: 10.1038/ejhg.2015.273

8. Tartaglia M, Gelb BD (2005) Noonan syndrome and related disorders: genetics and pathogenesis. Annu Rev Genomics Hum Genet 6:45–68. doi: 10.1146/annurev.genom.6.080604.162305

9. Cai W, Carlson SW, Brelsfoard JM, Mannon CE, Moncman CL, Saatman KE, Andres DA (2012) Rit GTPase signaling promotes immature hippocampal neuronal survival. J Neurosci 32 (29):9887–9897. doi: 10.1523/JNEUROSCI.0375-12.2012

10. Mir S, Cai W, Andres DA (2017) RIT1 GTPase Regulates Sox2 Transcriptional Activity and Hippocampal Neurogenesis. J Biol Chem 292 (6):2054–2064. doi: 10.1074/jbc.M116.749770

11. Tang Z, Li C, Kang B, Gao G, Li C, Zhang Z (2017) GEPIA: a web server for cancer and normal gene expression profiling and interactive analyses. Nucleic Acids Res 45 (W1):W98–W102. doi: 10.1093/nar/gkx247

12. Rhodes DR, Yu J, Shanker K, Deshpande N, Varambally R, Ghosh D, Barrette T, Pandey A, Chinnaiyan AM (2004) ONCOMINE: a cancer microarray database and integrated data-mining platform. Neoplasia 6 (1):1–6. doi: 10.1016/s1476-5586(04)80047-2

13. Shai R, Shi T, Kremen TJ, Horvath S, Liau LM, Cloughesy TF, Mischel PS, Nelson SF (2003) Gene expression profiling identifies molecular subtypes of gliomas. Oncogene 22 (31):4918–4923. doi: 10.1038/sj.onc.1206753

14. Diez-Villanueva A, Mallona I, Peinado MA (2015) Wanderer, an interactive viewer to explore DNA methylation and gene expression data in human cancer. Epigenetics Chromatin 8:22. doi: 10.1186/s13072-015-0014-8

15. Bredel M, Bredel C, Juric D, Harsh GR, Vogel H, Recht LD, Sikic BI (2005) Functional network analysis reveals extended gliomagenesis pathway maps and three novel MYC-interacting genes in human gliomas. Cancer Res 65 (19):8679–8689. doi: 10.1158/0008-5472.CAN-05-1204

16. Sun L, Hui AM, Su Q, Vortmeyer A, Kotliarov Y, Pastorino S, Passaniti A, Menon J, Walling J, Bailey R, Rosenblum M, Mikkelsen T, Fine HA (2006) Neuronal and glioma-derived stem cell factor induces angiogenesis within the brain. Cancer Cell 9 (4):287–300. doi: 10.1016/j.ccr.2006.03.003

17. Murat A, Migliavacca E, Gorlia T, Lambiv WL, Shay T, Hamou MF, de Tribolet N, Regli L, Wick W, Kouwenhoven MC, Hainfellner JA, Heppner FL, Dietrich PY, Zimmer Y, Cairncross JG, Janzer RC, Domany E, Delorenzi M, Stupp R, Hegi ME (2008) Stem cell-related “self-renewal” signature and high epidermal growth factor receptor expression associated with resistance to concomitant chemoradiotherapy in glioblastoma. J Clin Oncol 26 (18):3015–3024. doi: 10.1200/JCO.2007.15.7164

18. Thul PJ, Lindskog C (2018) The human protein atlas: A spatial map of the human proteome. Protein Sci 27 (1):233–244. doi: 10.1002/pro.3307

19. saba LPaJD (2014) Getting a Clue from 1q: Gain ofChromosome 1q in Cancer. Journal of Cancer Biology and Research 2 (3):5

20. Chou CH, Shrestha S, Yang CD, Chang NW, Lin YL, Liao KW, Huang WC, Sun TH, Tu SJ, Lee WH, Chiew MY, Tai CS, Wei TY, Tsai TR, Huang HT, Wang CY, Wu HY, Ho SY, Chen PR, Chuang CH, Hsieh PJ, Wu YS, Chen WL, Li MJ, Wu YC, Huang XY, Ng FL, Buddhakosai W, Huang PC, Lan KC, Huang CY, Weng SL, Cheng YN, Liang C, Hsu WL, Huang HD (2018) miRTarBase update 2018: a resource for experimentally validated microRNA-target interactions. Nucleic Acids Res 46 (D1):D296–D302. doi: 10.1093/nar/gkx1067

21. Yuan J, Xiao G, Peng G, Liu D, Wang Z, Liao Y, Liu Q, Wu M, Yuan X (2015) MiRNA-125a-5p inhibits glioblastoma cell proliferation and promotes cell differentiation by targeting TAZ. Biochem Biophys Res Commun 457 (2):171–176. doi: 10.1016/j.bbrc.2014.12.078

22. Li Z, Qian R, Zhang J, Shi X (2019) MiR-218-5p targets LHFPL3 to regulate proliferation, migration, and epithelial-mesenchymal transitions of human glioma cells. Biosci Rep 39 (3). doi: 10.1042/BSR20180879

23. Amir S, Simion C, Umeh-Garcia M, Krig S, Moss T, Carraway KL, 3rd, Sweeney C (2016) Regulation of the T-box transcription factor Tbx3 by the tumour suppressor microRNA-206 in breast cancer. Br J Cancer 114 (10):1125–1134. doi: 10.1038/bjc.2016.73

24. Rebe C, Vegran F, Berger H, Ghiringhelli F (2013) STAT3 activation: A key factor in tumor immunoescape. JAKSTAT 2 (1):e23010. doi: 10.4161/jkst.23010

25. Kim JE, Patel M, Ruzevick J, Jackson CM, Lim M (2014) STAT3 Activation in Glioblastoma: Biochemical and Therapeutic Implications. Cancers (Basel) 6 (1):376–395. doi: 10.3390/cancers6010376

26. de Souza CF, Sabedot TS, Malta TM, Stetson L, Morozova O, Sokolov A, Laird PW, Wiznerowicz M, Iavarone A, Snyder J, deCarvalho A, Sanborn Z, McDonald KL, Friedman WA, Tirapelli D, Poisson L, Mikkelsen T, Carlotti CG, Jr., Kalkanis S, Zenklusen J, Salama SR, Barnholtz-Sloan JS, Noushmehr H (2018) A Distinct DNA Methylation Shift in a Subset of Glioma CpG Island Methylator Phenotypes during Tumor Recurrence. Cell Rep 23 (2):637–651. doi: 10.1016/j.celrep.2018.03.107

27. Li Z, Chen Y, An T, Liu P, Zhu J, Yang H, Zhang W, Dong T, Jiang J, Zhang Y, Jiang M, Yang X (2019) Nuciferine inhibits the progression of glioblastoma by suppressing the SOX2-AKT/STAT3-Slug signaling pathway. J Exp Clin Cancer Res 38 (1):139. doi: 10.1186/s13046-019-1134-y

28. Hagey DW, Klum S, Kurtsdotter I, Zaouter C, Topcic D, Andersson O, Bergsland M, Muhr J (2018) SOX2 regulates common and specific stem cell features in the CNS and endoderm derived organs. PLoS Genet 14 (2):e1007224. doi: 10.1371/journal.pgen.1007224

29. Schmitz M, Temme A, Senner V, Ebner R, Schwind S, Stevanovic S, Wehner R, Schackert G, Schackert HK, Fussel M, Bachmann M, Rieber EP, Weigle B (2007) Identification of SOX2 as a novel glioma-associated antigen and potential target for T cell-based immunotherapy. Br J Cancer 96 (8):1293–1301. doi: 10.1038/sj.bjc.6603696

30. Song WS, Yang YP, Huang CS, Lu KH, Liu WH, Wu WW, Lee YY, Lo WL, Lee SD, Chen YW, Huang PI, Chen MT (2016) Sox2, a stemness gene, regulates tumor-initiating and drug-resistant properties in CD133-positive glioblastoma stem cells. J Chin Med Assoc 79 (10):538–545. doi: 10.1016/j.jcma.2016.03.010

